# Evidence for secondary-variant genetic burden and non-random distribution across biological modules in a recessive ciliopathy

**DOI:** 10.1101/362707

**Authors:** Maria Kousi, Onuralp Söylemez, Aysegül Ozanturk, Sebastian Akle, Irwin Jungreis, Jean Muller, Christopher A. Cassa, Harrison Brand, Jill Anne Mokry, Maxim Y. Wolf, Azita Sadeghpour, Kelsey McFadden, Richard A. Lewis, Michael E. Talkowski, Hélène Dollfus, Manolis Kellis, Erica E. Davis, Shamil R. Sunyaev, Nicholas Katsanis

## Abstract

The influence of genetic background on driver mutations is well established; however, the mechanisms by which the background interacts with Mendelian loci remains unclear. We performed a systematic secondary-variant burden analysis of two independent Bardet-Biedl syndrome (BBS) cohorts with known recessive biallelic pathogenic mutations in one of 17 BBS genes for each individual. We observed a significant enrichment of *trans*-acting rare nonsynonymous secondary variants compared to either population controls or to a cohort of individuals with a non-BBS diagnosis and recessive variants in the same gene set. Strikingly, we found a significant over-representation of secondary alleles in chaperonin-encoding genes, a finding corroborated by the observation of epistatic interactions involving this complex *in vivo*. These data indicate a complex genetic architecture for BBS that informs the biological properties of disease modules and presents a model paradigm for secondary-variant burden analysis in recessive disorders.

---

A persistent hurdle in interrogating the role of genetic background in human genetic disorders is our limited understanding of the properties and distribution of contributory alleles. The challenge is particularly acute in rare disorders, in which the allele frequency of both causal variants and secondary contributory alleles (i.e. alleles in loci other than the primary locus) is often low; as such population-based studies are hampered by the lack of statistical power. At the same time, transitioning from a single-gene-centric to a systems-based disease architecture defined by biological modules can inform causality, penetrance and expressivity^1^.

Bardet-Biedl syndrome, a model ciliopathy, represents an opportunity to study secondary-variant burden. We and others have shown previously that BBS patients can carry secondary pathogenic variants in known BBS genes^2^. In rare examples, such alleles can modify penetrance^3^, whereas, more commonly, they are thought to modulate expressivity^4,5^. However, initial population-based studies have failed to detect an enrichment for secondary alleles in *trans* (i.e. alleles in loci other than the primary locus), suggesting that either some of the examples were exceptions, or that the incidence, distribution, and frequency of such alleles might be different than assumed *a priori*^6^. Here, we studied two BBS cohorts with unambiguous recessive pathogenic mutations in 17 established BBS genes to measure a) whether there is enrichment for secondary variants beyond the driver locus; and b) if so, whether the excess variation is concentrated within discrete disease modules or whether it is randomly distributed.

As a first step, we used targeted exon capture to sequence 102 families of Northern European^7^ ancestry (Discovery cohort), all of whom have *bona fide* pathogenic recessive mutations in one of 17 known BBS genes (*BBS1, BBS2, ARL6/BBS3, BBS4, BBS5, MKKS*/*BBS6, BBS7, TTC8/BBS8, BBS9, BBS10, TRIM32/BBS11, BBS12, MKS1/BBS13, CEP290*/*BBS14, WDPCP*/*BBS15, SDCCAG8/BBS16*, and *NPHP1*). We also sequenced an ethnically matched control cohort of 384 individuals (NEU control cohort) using the same exon capture technology (capture library, sequencing platform, and base-calling software).

Our base-calling method re-detected all 152 previously-identified pathogenic recessive mutations in our cases (50 families with compound heterozygous and 52 with homozygous mutations; **Supplementary Table S1**), indicating a negligible false negative rate. To eliminate the possibility of signal being driven by false positive events, we used Sanger sequencing to test all heterozygous burden-contributing secondary alleles at MAF<0.1% in both cases and controls. We confirmed 21 variants in the Discovery cohort and 45 in the NEU control cohort.

We then asked whether individuals with a clinical diagnosis of BBS have an increased burden of *trans* secondary variants that lie beyond the primary locus, in addition to the recessive diagnostic changes. If the recessive event at the primary locus is sufficient for disease manifestation (null hypothesis), we would expect no difference in allele burden between BBS patients and controls; however, if additional variants in *trans* with the primary locus contribute to disease, we would expect to see an enrichment of such changes in cases (burden hypothesis).

To distinguish between the two posits, we used a collapsing and combine (CMC) test^8^ of rare variants at varying in-cohort MAF cutoffs, restricting our analysis to nonsynonymous heterozygous variants beyond the primary locus. Within NEU descent individuals (n=84 cases and n=384 controls), we observed a significant enrichment for burden-contributing heterozygous *trans* variants beyond the primary driver locus (**Table 1**) in cases versus controls (CMC test p=8×10^−3^; OR=1.77; **Fig. 1A**) at MAF=0.1%. This result also withheld when evaluating the entire discovery cohort (n=102 cases and n=384 controls), by including 18 additional individuals of mixed European ethnicity (p=0.008; OR=2.12 at MAF=0.9%). Also supporting the burden hypothesis, BBS cases (n=102) showed a significant excess of singletons (alleles found just once in any cohort) versus controls in the distribution of non-synonymous variants beyond the “driver” locus (p=0.001; OR=2.08; **Fig. 1A** and **Table S1**). Doubletons and tripletons were found to occur equally likely between cases and controls, arhuing against significant population substructure affecting the observed signal (**Table S1**).

**Table 1.** Burden analysis for variants beyond the primary driver locus.

| Discovery cohort |  |  |  |  | Replication cohort |  |  |  |  |
| --- | --- | --- | --- | --- | --- | --- | --- | --- | --- |
| Cases | Controls | OR<sup>a</sup> | P<sup>a</sup> | MAF cutoff<sup>b</sup> | Cases | Controls | OR | P | MAF cutoff |
| 102 | 384 | 1.77 | $8 \times 10^{-3}$ | 0.9% | 89 | 466 | 2.11 | $1 \times 10^{-3}$ | 0.2% |
| 84<sup>c</sup> | 384 | 2.12 | $8 \times 10^{-3}$ | 0.1% | | | | | |

<sup>a</sup> Odds ratio (OR) and one-sided p-value computed using Fisher's Exact test at the optimal in-cohort frequency threshold <sup>b</sup> Optimal frequency threshold selected using a variable threshold approach to incorporate allele frequency information into burden analysis. Significance of the threshold test was evaluated using 10000 permutations at alpha=0.05. <sup>c</sup> Subset of individuals with Northern European ancestry.

**Figure 1.**
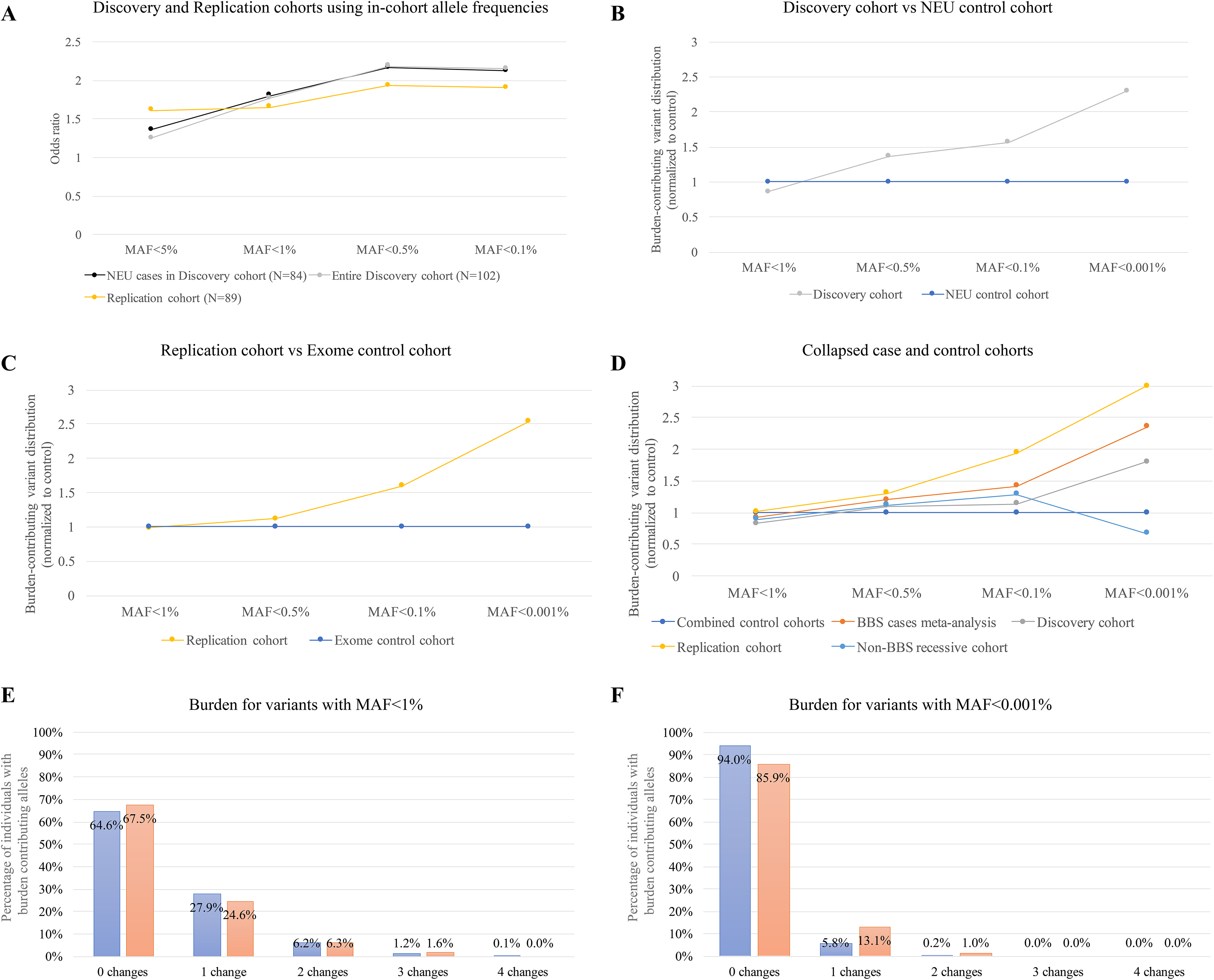
Distribution of burden-contributing variation across case and control cohorts. **(A)** Collapsing and combine (CMC) test of rare non-synonymous heterozygous variants outside the primary driver locus at varying in-cohort minor allele frequency (MAF) cutoffs; **(B)** Values of burden-contributing variation between BBS cases of the Discovery cohort (N=102) and the cohort of Northern European controls (N=384); **(C)** BBS cases of the Replication cohort (N=89) and the Exome control cohort (N=466); and **(D)** collapsed cohorts, across four MAF bins (1%, 0.5%, 0.1% and 0.001%). **(A)** The Discovery case cohort shows a 2.3-fold enrichment for ultra-rare (MAF<0.001%) alleles compared to controls, **(B)** the Replication cohort shows a 2.5-fold enrichment of such alleles compared to the exome control cohort and **(C)** the BBS case meta-analysis a 2.4-fold enrichment compared to the combined control cohorts. **(E)** Distribution of individuals with burden-contributing alleles with MAF<1%; and **(F)** with MAF<0.001% in control individuals (blue bars) and BBS cases (orange bars).

As a further test, we next evaluated the distribution of synonymous changes across the case-control cohorts. In the absence of population structure, synonymous variants should be distributed evenly among cases and controls; we found no significant difference in the distribution of singleton synonymous variants in the entire Discovery cohort (n=102), suggesting that population structure or another technical artifact, is not a confounder (**Table S1**). Likewise, no differences were observed in the distribution of doubletons or tripletons (**Tables S1 and S2**).

We next classified variants according to their population frequency in ExAC, reasoning that alleles that are rare in the population are more likely to represent disruptions of functional nucleotides; such an analysis is facilitated by the availability of large-scale reference datasets of human genetic variation that lend the opportunity to recognize low-frequency genetic variants (on the order of the disease prevalence), rather than being limited to the in-cohort level allelic frequency. For each BBS case with a recessive primary locus, we measured the incidence of burden alleles in the other 16 BBS genes at four ExAC-based MAF cutoffs: 1%, 0.5%, 0.1%, and 0.001%. Consistent with the CMC test, we found a significant enrichment for MAF<0.001% (“ultra-rare alleles”) in cases versus controls (2.3-fold enrichment; p=0.03; OR=2.46; **Fig. 1B**).

To confirm our findings, we assembled a replication cohort of 555 individuals composed of 89 cases with a secured molecular genetic BBS diagnosis in one of the 17 BBS genes (Replication cohort), and 466 control individuals (Exome control cohort). Using the CMC test, we once again observed a significant enrichment for burden alleles in cases versus controls (p=0.001; OR=2.11 at MAF=0.2%; **Fig. 1A**), similar to the discovery case-control dataset. Using population-based allele frequencies, we observed a significant (2.5-fold) enrichment of ultra-rare alleles (MAF<0.001%; 17 changes / 89 cases versus 35 changes / 466 controls; p=2×10^−3^, OR=2.88; **Fig. 1C**), once again similar to the Discovery cohort. Critically, testing for burden in synonymous variants showed no association (p=0.21), while testing for balancing of singletons, doubletons and tripletons was likewise bereft of evidence of population stratification bias (p=0.21 for singletons, p=0.33 for doubletons and p=0.42 for tripletons; **Table S3**); we attribute this observation to the stringent filtering of population-based MAFs (always considering the highest possible MAF) that reduced the possible inflation of population-specific alleles.

Meta-analysis of the two case cohorts (Discovery and Replication) using a Cochran-Mantel-Haenszel (CMH) procedure corroborated the enrichment of rare secondary burden alleles in BBS patients compared to controls taking into account the cohort status as a stratification variable (p< 0.00001; CMH pooled OR=1.94). The enrichment withstood when considering population-based allele frequencies in BBS (29 changes / 191 cases compared to 53 changes / 850 control individuals p=4×10^−4^; OR=2.58; **Fig. 1D**). To obtain further evidence of the specific MAF fraction contributing to burden, we counted and subsequently normalized the number of individuals with changes in each of the four established MAF bins (1%<MAF<0.5%, 0.5%<MAF<0.1%, 0.1%<MAF<0.001%, and 0.001%<MAF<0%). Consistent with our previous observations, the signal for burden was driven exclusively by ultra-rare alleles (0.001%<MAF<0%; p=0.0004; OR=2.58; **Supplementary Fig. 1**).

Notably, we found that *trans*-acting secondary variants in BBS genes were more likely to disrupt strongly-conserved genomic segments in the combined BBS cases than in combined controls. We focused on Synonymous Constraint Elements (SCEs) in 29 mammalian genomes, which show protein-coding conservation and also strong non-coding constraint in synonymous positions, and thus are likely to contain overlapping functional elements, such as RNA secondary structures, or binding sites for regulatory proteins^9^. We found that secondary variants overlap SCEs for 5/76 BBS case variants (6.6%) but only for 6/371 control variants (1.6%) (hypergeometric p=0.025). Though these numbers are low and as such warrant caution, our data do suggest that burden-contributing variants are more likely to disrupt functional elements in BBS cases.

Next, to gain further insight into the properties that distinguish the exomes of BBS-diagnosed individuals from the exomes of individuals who carry “chance” recessive changes in BBS genes, we parsed ~10,000 exomes from the Baylor Molecular Diagnostics laboratory; we identified 50 individuals with a clinical genetic disorder other than BBS who carry homozygous rare (MAF<1%) alleles in one of the BBS genes (“non-BBS recessive” cohort). In contrast to our 191 *bona fide* BBS cases, the 50 “non-BBS recessive” individuals showed no significant burden in ultra-rare BBS alleles compared to the 850 combined control cohorts (p=0.76; **Fig. 1D**), consistent with the model that burden alleles contribute to the clinical manifestation of BBS.

In addition to burden alleles, we also studied differences in the primary locus mutations of BBS cases and “non-BBS recessive” individuals. Overall, 37% (71/191) of the BBS patients harbor two highly disruptive (nonsense, frameshifts, deletions, etc) mutations in the primary locus in contrast to only 2% (1/50) of the “non-BBS recessive” cases (p=4×10^−8^). Furthermore, for the individuals harboring at least one missense mutation in the primary locus, we attributed a weight for the impact of these nonsynonymous changes using BLOSUM (score ranges between −3 to 3, with lower scores denoting more deleterious substitutions)^10^. BLOSUM scores of the BBS case variants were significantly lower and thus likely more disruptive than the scores for the “non-BBS recessive” individuals, with 76% (90/118) of the *bona fide* BBS cases having a negative BLOSUM score, as opposed to only 43% (21/49) of the “non-BBS recessive” individuals (p=4×10^−5^; **Supplementary Figure 2**). Taken together, our data are consistent with increased mutational load both in the primary locus and in secondary *trans*-acting variants that may interact epistatically with the primary locus.

Next, we turned our attention to the question of whether the observed burden alleles are distributed randomly across the BBS genes. For this question, we tested the observed versus expected probability of an individual with a mutation in any given BBS gene to have a secondary variant in a second BBS gene. We found enrichment for a subset of genes. For example, if the primary recessive mutation occurred in *BBS5*, the secondary mutation was 13-fold more likely to occur in *BBS2* than in other BBS genes (*BBS5* → *BBS2*; p=0.0005). Though the small number of interactions merit caution, we observed another three such pairings to be significantly enriched: *BBS9* → *BBS10* (10-fold, p=0.0014); *BBS12* → *BBS4* (13-fold, p=0.0005); *SDCCAG8* → *BBS1* (26-fold, p=0.0019) (**Fig. 2A**). By contrast, *BBS1* and *BBS10*, which contribute almost 40% of the recessive drivers in our cohorts, were not enriched for secondary *trans* variants in any other specific BBS gene, suggesting that the frequency of primary drivers is not a predictor of interactions. Neither our burden observations nor the specific pairings were driven by gene size; for example, *CEP290* was the largest of the evaluated BBS loci, but showed neither a mutational enrichment nor an interaction enrichment in our analyses.

**Figure 2.**
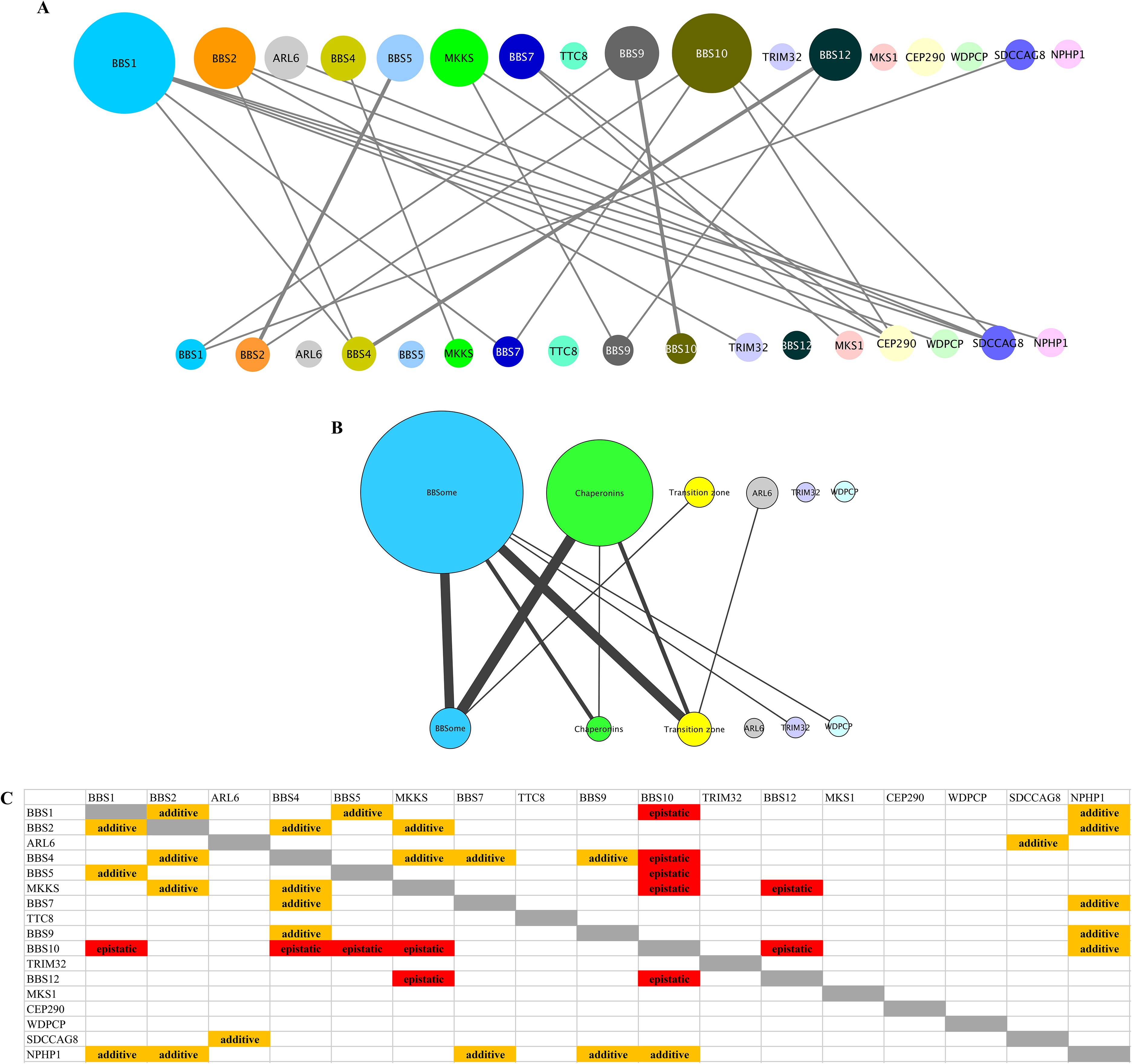
Genetic and modular interactions in BBS cases and controls. In each panel the primary loci are represented as circles in the top half and the loci contributing to mutational load as circles in the bottom half of each panel respectively. The size of the circles corresponds to the number of individuals carrying changes in the primary and burden contributing loci, respectively and the thickness of the lines connecting primary and burden-contributing loci corresponds to the frequency at which a genetic interaction is observed with thicker lines representing more common interactions. **(A)** In the meta-analysis of BBS cases *BBS1, BBS2, BBS10* and *BBS12* harbor recessive driver alleles most frequently but there is an even distribution of burden-contributing alleles across the 17 BBS loci. **(B)** Analysis of modules within the BBS proteome (BBSome: *BBS1, BBS2, BBS4, BBS5, BBS7, TTC8*, and *BBS9*; Chaperonin complex: *MKKS, BBS10*, and *BBS12*; Transition zone: *MKS1, CEP290, SDCCAG8*, and *NPHP1* and the three genes that belong to no defined yet module: *ARL6*; *TRIM32*; *WDPCP*) revealed that the components of the chaperonin complex are driving the majority of modular interactions in BBS. (**C**) Schematic showing the functional outcome (epistatic versus additive) upon suppression of select modular interactions using an *in vivo* zebrafish system. Epistatic interactions among loci are shown in red and additive genetic interactions are shown in orange.

To probe deeper into the nature and distribution of interactions, we wondered whether this burden is distributed randomly across the 17-BBS gene set. We noted that primary-secondary locus interactions extend beyond the observed pairwise relationships to macromolecular complexes. We were struck that two of the four enriched interaction pairings include a chaperonin-encoding gene (*BBS10* and *BBS12*). We therefore clustered the 17 BBS genes into three previously-defined modules: the BBSome complex (BBS1, BBS2, BBS4, BBS5, BBS7, TTC8 and BBS9)^11-13^; the transition zone complex (MKS1, CEP290, SDCCAG8, and NPHP1)^7,14,15^, and the chaperonin complex (MKKS, BBS10 and BBS12)^16,17^. For the remaining three genes/proteins (WDPCP, TRIM32, and ARL6), we have no information about their macromolecular complexes. Restricting ourselves to alleles at MAF<0.001% (the bin with the most robust evidence for burden enrichment) we collapsed all alleles per module and plotted all observed interactions. We then calculated the number of interactions within and across modules (**Fig. 2B**). The distribution was non-random: across the combined BBS patient set of 191 individuals, most recessive events were contributed by chaperonin-encoding components (p=4×10^−3^; **Fig. 2B**), suggesting that this module potentially has the most physiologically potent interaction capability.

As part of our gene discovery work, we have tested pairs of BBS genes for their ability to interact *in vivo*. In this paradigm, we have suppressed each gene either alone or in pairs in sub-effective doses in zebrafish embryos and asked whether double suppression affected the severity of the phenotype in an additive or multiplicative fashion^18^. *Post-hoc* retrieval of interaction data across 19 gene pairs revealed 13 additive and six epistatic interactions (**Fig. 2C**)^17,19-22^. Strikingly, all six observed epistatic interactions involved one of the chaperonins, whereas none of the non-chaperonin pairings showed epistasis (p=0.003).

In this study, we provide direct evidence for the enrichments of ultra-rare alleles in BBS patients with diagnosed recessive mutations. The degree of enrichment (~2.5-fold) is similar in both our cohorts and is only detectable for ultra-rare alleles (MAF<0.001%), consistent with population-level selection against these mutations, and indicating that they are likely deleterious and occurring at functional positions. We suspect that this occurs because the majority of variants are likely detrimental to protein function at this MAF; the study of larger cohorts should allow for the detection of more common alleles, especially if it is coupled to rigorous functional studies to determine their effect. We also note that the non-random distribution of these variants underscores biological substructure in this module. Although we are cautious to avoid over-interpreting our data, our observations support the intuitive expectation that genetic interactions within macromolecular complexes are more likely to be additive, but interactions between complexes are more likely to be epistatic. For BBS, understanding the biochemical interplay between the chaperonin module and the BBSome will likely improve further our understanding of the role of the additional variation.

We do not know the contribution of burden to the phenotype. If the discovered variants were necessary for classically-defined penetrance, we would anticipate a substantial fraction of asymptomatic individuals with recessive BBS mutations, which is not true. At the same time, pure modifiers of expressivity would have no mechanistic reason to be enriched in our cohort, since we did not pre-select for endophenotypes. The most parsimonious explanation is that recessive mutations are necessary to cause disease but are not sufficient to meet the threshold of a clinical BBS diagnosis; that is achieved through additive or multiplicative interactions with deleterious variants in other components of the module. Our current data indicate that *a*pproximately 22% of our patients carry one or more additional ultra-rare variants at the MAF<0.1% cutoff (at which significant enrichment for secondary *trans* variants is detected with the CMC test analysis). This is likely to be an under-estimate of burden since additional *trans* alleles are likely to be a) of higher frequency than we have statistical power to detect in this cohort; b) lie elsewhere in the known BBS genes; and c) are likely to also exist in non-BBS-causing recessive loci that are nonetheless relevant to the function of the BBS biological modules.

Finally, we emphasize the fact that studies of genetic burden in rare genetic disorders can be powered significantly by biological insight. In our datasets, had we attempted agnostic genome-wide tests, the observed burden would have been invisible because of the size of the denominator. Focusing on comprehensive set of known disease-causing genes allows a sensible, functionally-driven hypothesis to be tested that also avoids the well-established pitfalls of single-gene candidate studies. Indeed, our findings are concordant with recent studies of other genetic disorders, including Charcot-Marie-Tooth disease^24^ and retinitis pigmentosa^25^. We speculate that the systematic measurement of burden of other genetically-heterogeneous disorders in which all the causal genes are considered as one “baseline” entity (essentially a functional operon) will reveal similar burden observations, which in turn can be used to understand biological and biochemical substructure.

## Methods

### Research participants

We assessed two cohorts of individuals who fulfilled the clinical diagnostic criteria for BBS^26^. The Discovery cohort comprised 102 patients of northern European descent and the Replication cohort 89 unrelated individuals of mixed ethnicity, respectively (**Supplementary Table S4**). The control cohorts included 384 northern European individuals (NEU control cohort) and 466 whole exome sequenced unaffected individuals of mixed ethnicity (Exome control cohort). A cohort of 50 individuals with a non-BBS genetically identified disorder, who were harboring at least two alleles with MAF<1% in one of the 17 BBS loci evaluated, was also assembled (“non-BBS recessive” cohort). Informed consent was obtained from all controls, BBS cases, and their willing family members according to protocols approved by the Institutional Review Boards of the Duke University Medical Center, the Université de Strasbourg (“Comité Protection des Personnes” EST IV, N°DC-20142222) and the Baylor College of Medicine. We obtained peripheral whole blood samples from participants and extracted DNA according to standard methods.

### Targeted exome capture and next generation sequencing

The BBS patients and controls were sequenced either by regional capture of the exons and intron-exon boundaries of 785 ciliary genes prioritized from the ciliary proteome^27^, or by direct Sanger sequencing of each exon in a CLIA laboratory^28^. We captured regions of interest with a custom NimbleGen targeted liquid capture (12,000 exons; 1.9 Mb of target) according to manufacturer’s instructions, pooled 23 samples/pool, and subsequently performed next-generation sequencing with an Illumina HiSeq2000 platform (paired-end 100bp reads with two pools/lane; 10 lanes total). The samples were sequenced with a mean of 110x coverage with 92% of bases achieving >20x coverage and 99% of targeted regions captured. All discovered alleles at MAF<1% were confirmed by secondary Sanger sequencing. The “non-BBS recessive” cohort was sequenced by whole exome capture methodology and underwent the same analysis as previously described^29^.

### Mutation burden analysis

We performed a unidirectional gene-based association test^8^ using the mutational target of 17 BBS genes as a single grouping unit. To assess the contribution of burden alleles beyond the primer driver locus, each BBS patient’s genotype at the respective driver locus was set to homozygous reference. Missing genotypes in both cases and controls were imputed to reference. Restricting to nonsynonymous single nucleotide variants with high impact on protein function (missense and nonsense consequences as annotated by SnpEff v4.3)^30^, we performed collapsing and combine (CMC) test of rare variants at various in-cohort minor allele frequency (MAF) cutoffs as well as at the optimal in-cohort frequency cutoff selected using a variable threshold (VT) approach^31^, as implemented in RVTESTS package^32^. To mitigate potential bias due to imperfect ethnicity matching between cases and controls, we repeated the burden analysis excluding 18 individuals with non-Northern European ancestry.

We assessed mutational burden in BBS cases, controls and the cohort of “non-BBS recessive” individuals with a known primary recessive driver gene (**Supplementary Table S4**). For inclusion, variants fulfilled the following criteria: 1) likely disruptive to protein sequence i.e. nonsynonymous, nonsense, frameshifting, *bona fide* splice sites within three base pairs of exonintron junctions, and CNVs described previously^19,20^; 2) ethnically-matched MAF<1% (as obtained from the ExAC browser: http://exac.broadinstitute.org/ and/or the Greater Middle East (GME) Variome: http://igm.ucsd.edu/gme/index.php) and 3) restricted to the 17 BBS genes sequenced across all three cohorts. The statistical significance between control and affected individuals with variants likely contributing to burden was compared among groups (Discovery cohort vs NEU control cohort; Replication cohort vs Exome control cohort; Combined BBS cohorts (Discovery+Replication) vs Combined control cohorts (NEU control cohort+ Exome control cohort) with a two-tailed Fisher’s exact probability test.

### Synonymous Constraint Elements

We searched for Synonymous Constraint Elements (SCEs) in every coding sequence in CCDS Homo Sapiens release 9 using FRESCo software^33^. FRESCo analysis was performed using a window size of 9 and sequences and distances taken from the 29 mammal alignment^34^. The resulting list of SCEs is available at https://data.broadinstitute.org/compbio1/SynonymousConstraintTracks/ESS.hg19.bed.gz. We tested each burden SNV in each individual and found the following changes to be contained in an SCE: for cases: AR201-06, BBS9: p.Ser753Phe; AR831-03, BBS4: p.Pro13Ser; KK011-04, SDCCAG8: p.Glu532Lys; Fam93_NNMR18, BBS4: p.R295Q; Fam14_AKX44, SDCCAG8: p.Q505E; for controls: 37 - 10101398_C31NDACXX-7-IDMB39, NPHP1: p.S29F; 246 - 10104594_C30WYACXX-4-IDMB66, BBS9: p.S788F; DM414-1000, NPHP1: p.M616I; DM773-1000, NPHP1: p.I45L; DM1377-1000, ARL6: p.G72R; DM1377-1001, BBS9: p.S788F.

### BLOSUM scoring

BLOSUM score was calculated using the blosum62 matrix obtained from biopython version 1.58. The BLOSUM score ranges from −3 to 3 with lower numbers indicating biochemically more different pairs of amino acids and higher scores indicating more-radical amino acid changes. For BBS cases with two missense changes at the recessive locus, the change with the higher BLOSUM score was taken into consideration.

## Acknowledgements

We thank the patients and their families for participating in this study, as well as their caring physicians that provided clinical information and referred the patients for diagnostic analyses. We thank Anne-Sophie Jaeger and Manuela Antin for the technical assistance. This study was supported by the NIH grants GM121317-13, HD042601, and DK072301 (N.K.). R01 HG004037 (I.J.) and GENCODE Wellcome Trust grant U41 HG007234 (I.J.). R.A.L. is a senior scientific investigator of research to prevent blindness. N.K. is a distinguished Jean and George Brumley Professor.

## Author contributions

M.K., E.E.D. S.S. and N.K. designed the overall study. M.K., A.O., and K.M. performed the experimental work by genotyping and analyzing the sequencing data. O.S., S.A., C.A.C., M.K., H.R. and M.E.T., and S.S. performed the statistical analysis assessing mutational burden. I.J., M.Y.W. and M.K. performed the analyses assessing the impact of the identified changes on genomic sequence signatures. J.M., J.A.M. R.A.L. and H.D. contributed samples that were analyzed in this study. A.S. collated the genetic information of the Discovery cohort. M.K. performed the collective analysis of genetic data and modular interactions. All authors have read and commented on the manuscript that was written by N.K. and M.K.

---

**Supplementary Fig. 1. Distribution of burden-contributing variation across case and control cohorts in each of four discrete MAF bins. (A)** Values of burden-contributing variation between BBS cases of the Discovery cohort (N=102) and the cohort of Northern European controls (N=384); **(B)** BBS cases of the Replication cohort (N=89) and the Exome control cohort (N=466); **(C)** BBS cases of the meta-analysis (N=191) and the combined control cohorts (N=850); and **(D)** collapsed cohorts, across four MAF bins (1%<MAF<0.5%, 0.5%<MAF<0.1%, 0.1%<MAF<0.001%, and 0.001%<MAF<0%). **(A)** The Discovery case cohort shows a 2.3-fold enrichment for ultra-rare (0.001%<MAF<0%) alleles compared to controls, **(B)** the Replication cohort shows a 2.5-fold enrichment of such alleles compared to the exome control cohort and **(C)** the BBS case meta-analysis a 2.4-fold enrichment compared to the combined control cohorts.

**Supplementary Fig. 2.** Estimate of protein impact for the least disruptive of the diagnostic variants for BBS cases and non-BBS recessive individuals, with at least one missense change in the primary locus. With high BLOSUM62 scores denoting biochemically similar amino acid changes and lower scores marking radical amino acid changes, the graph shows evidence for *bona fide* BBS cases harboring more disruptive variants.

**Table S1.**
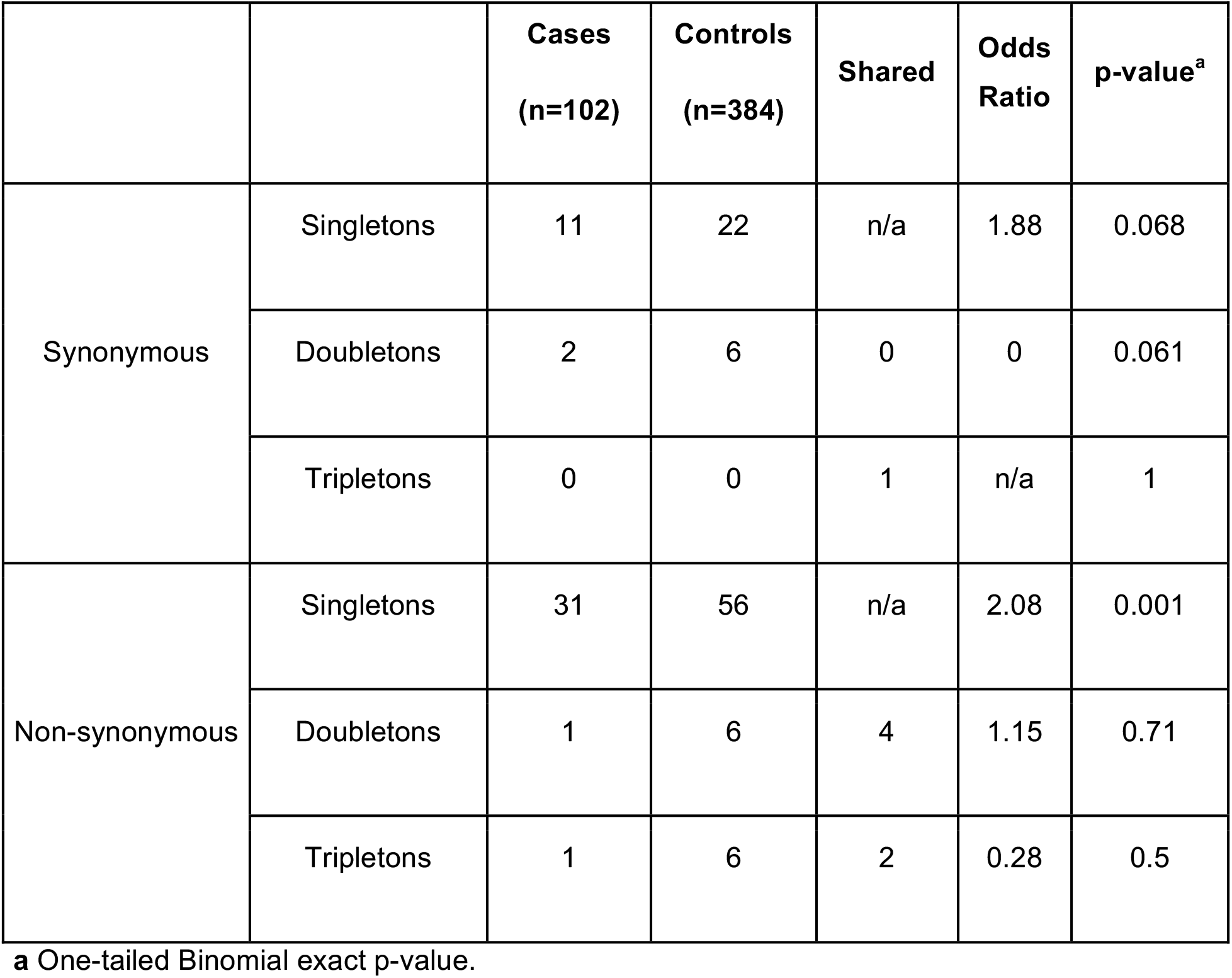
Distribution of synonymous and non-synonymous singletons, doubletons and tripletons between cases and controls in the original discovery cohort.

|  |  | **Cases**<br>**(n=102)** | **Controls**<br>**(n=384)** | **Shared** | **Odds**<br>**Ratio** | **p-value<sup>a</sup>** |
| --- | --- | --- | --- | --- | --- | --- |
| Synonymous | Singletons | 11 | 22 | n/a | 1.88 | 0.068 |
|  | Doubletons | 2 | 6 | 0 | 0 | 0.061 |
|  | Tripletons | 0 | 0 | 1 | n/a | 1 |
| Non-synonymous | Singletons | 31 | 56 | n/a | 2.08 | 0.001 |
|  | Doubletons | 1 | 6 | 4 | 1.15 | 0.71 |
|  | Tripletons | 1 | 6 | 2 | 0.28 | 0.5 |

**a** One-tailed Binomial exact p-value.

**Table S2.** Distribution of synonymous and non-synonymous singletons, doubletons and tripletons between cases and controls when cases with non-Caucasian ancestry were excluded.

|  |  | **Cases**<br>**(n=84)** | **Controls**<br>**(n=384)** | **Shared** | **Odds**<br>**Ratio** | **p-value<sup>a</sup>** |
| --- | --- | --- | --- | --- | --- | --- |
| Synonymous | Singletons | 8 | 22 | n/a | 1.66 | 0.15 |
|  | Doubletons | 0 | 6 | 0 | 0 | 0.12 |
|  | Tripletons | 0 | 0 | 1 | n/a | 1 |
| Non-synonymous | Singletons | 26 | 61 | n/a | 1.95 | 0.004 |
|  | Doubletons | 1 | 8 | 2 | 0.45 | 0.32 |
|  | Tripletons | 1 | 6 | 2 | 0.23 | 0.16 |

**a** One-tailed Binomial exact p-value.

**Table S3.** Distribution of synonymous and non-synonymous singletons, doubletons and tripletons between cases and controls in the replication cohort.

|  |  | **Cases**<br>**(n=89)** | **Controls**<br>**(n=466)** | **Shared** | **Odds**<br>**Ratio** | **p-value<sup>a</sup>** |
| --- | --- | --- | --- | --- | --- | --- |
| Synonymous | Singletons | 14 | 55 | n/a | 1.33 | 0.21 |
|  | Doubletons | 4 | 6 | 2 | 0.4 | 0.33 |
|  | Tripletons | 0 | 4 | 4 | 0.68 | 0.42 |
| Non-synonymous | Singletons | 30 | 116 | n/a | 1.35 | 0.038 |
|  | Doubletons | 4 | 19 | 4 | 0.47 | 0.11 |
|  | Tripletons | 0 | 1 | 7 | 4.74 | 0.98 |

**a** One-tailed Binomial exact p-value.

